# Axonal tree morphology and signal propagation dynamics improve interneuron classification

**DOI:** 10.1101/414615

**Authors:** Netanel Ofer, Orit Shefi, Gur Yaari

## Abstract

Neurons are diverse and can be differentiated by their morphological, electrophysiological, and molecular properties. Current morphology-based classification approaches largely rely on the dendritic tree structure or on the overall axonal projection layout. Here, we use data from public databases of neuronal reconstructions and membrane properties to study the characteristics of the axonal and dendritic trees for interneuron classification. We show that combining signal propagation patterns observed by biophysical simulations of the activity along ramified axonal trees with morphological parameters of the axonal and dendritic trees, significantly improve classification results compared to previous approaches. The classification schemes introduced here can be utilized to robustly classify neuronal subtypes in a functionally relevant manner. Our work paves the way for understanding and utilizing form-function principles in realistic neuronal reconstructions.

## Introduction

Quantitative analysis of neuronal types and their properties is critical for better understanding and deciphering brain function (*1, 2*). Despite the attempts to standardize the terminology for neuronal types, there is no clear consensus regarding neuron nomenclature, leaving neuronal classification as an ongoing challenge (*3, 4*). To date, interneuron classification is based on morphology (*5*), membrane and firing patterns (*6, 7*), connectivity patterns (*8*), neurochemical markers (*9*), transcriptome (*10–12*), and epigenomics (*13*). The morphology-based classification approaches include dendritic tree geometry (*14, 15*) and axonal projection (*16, 17*), where directionalities of axons are taken into account. The interneuron’s axonal tree arbor enables better classification of cell types than the dendritic tree (*8, 18*). Topological persistence-based methods were also developed to support comparisons between individual neurons and classification of neurons (*19, 20*). Topological motifs of the axonal tree were found to differentiate interneurons and pyramidal cells (*21, 22*). So far, no studies have used the geometrical properties of the axonal tree, specifically the axonal branch diameters and lengths, for neuronal classification.

Different types of neurons have different ion channels with various kinematics and densities, spreading across the soma, axons, and dendrites (*23, 24*). As part of the Blue *Brain Project* (BBP), evolutionary algorithms were used to fit the experimental recordings of rat cortical neurons with specific ion channel types and parameters. Firing patterns are commonly defined by neuronal responses to step currents at the soma. Combinations of continuous, delayed, and bursting onset patterns, with accommodating, non-accommodating, stuttering, irregular, and adapting steady-state behaviors, led to establishing eleven electrical types (e-types), ten of which exist in interneurons and one in pyramidal cells. The distribution of each of the ion channels along specific neuronal types and cortical layers as well as the fitted parameters are indicated in the Neocortical Microcircuit Collaboration Portal (NMC) (*25*). Hence, activity-based neuronal classification is a promising and interesting path that remains to be explored.

Here, we have leveraged the advancement of imaging techniques that led to growth in high-resolution 3D reconstructions along with the development of big neuronal morphology databases, such as the *Blue Brain Project* (*26*), the Allen *Institute Brain Atlas* (*27*), and *NeuroMorpho.Org* (*28*), to classify interneurons into subtypes based on their morphology and activity. We first classified interneurons based on axonal tree morphology parameters, obtaining fairly accurate discrimination. Adding dendritic tree morphology to the axonal one improved the prediction rates. Finally, we considered an axonal tree activity-based neuronal classification and further improved the classification’s results. Building a classification scheme based on all these features is shown here to robustly classify neurons in a functionally relevant manner.

## Results

### Classification of interneuron types by morphology

To classify interneurons based on axonal tree morphology, high-resolution traced neurons were analyzed. For this purpose, neuron reconstructions were downloaded from the *NeuroMorpho.Org* database, and filtered for several criteria to obtain a high-quality dataset for classification (Table 1). Only neurons from a cortex with at least 10 axonal branches and 1,000 axonal segments were included. To achieve high precision in axonal tree geometry, only neurons with at least 10 axonal diameter values measured were included. The resulting filtered dataset is diverse because the interneurons were taken from different cortical layers of male and female rats (n=312, 78%) and mice (n=90, 22%), and were analyzed by different labs. We focus here on the most prominent interneuron types: basket cell (BC), Martinotti cell (MC), chandelier cell (CHC), neurogliaform cell (NGF), bitufted cell (BTC), double-bouquet cell (DBC), and bipolar cell (BP) (*29*). Figure 1A–G show representative examples of the interneuron types.

**Table 1:** Neuron reconstructions filtration. The table summarizes the number of neurons that were included after each filtration step according to their types.

| Filter criteria | basket | Martinotti | neurogliaform | bitufted | double bouquet | chandelier | bipolar |
| --- | --- | --- | --- | --- | --- | --- | --- |
| All data in NeuroMorpho.Org 7.4 | 829 | 294 | 209 | 93 | 64 | 68 | 606 |
| Cortex only | 564 | 228 | 139 | 88 | 63 | 36 | 40 |
| With both axon and dendrite data | 508 | 224 | 127 | 56 | 56 | 31 | 40 |
| $\geq 10$ axonal branches and $\geq 1,000$ axonal segments | 437 | 193 | 122 | 53 | 52 | 29 | 34 |
| $\geq 10$ axonal diameter values measured | 196 | 99 | 40 | 20 | 20 | 17 | 10 |

Each neuron reconstruction is characterized by 28 features that can be divided into three categories: overall topology, branch length, and diameter (Table 2). Overall topology measurements include the number of branches, branch order, Sholl analysis, the axonal tree size, and symmetry. Branch length-related parameters include the total length, branch lengths, path lengths, and the branch length divided by the square root of the diameter. The diameter-related parameters include the branch diameter and the geometric ratio (GR). GR is defined as the ratio between the sum of the diameter of the two daughter branches and the mother branch, to which a 3/2 power exponent is applied. These features are expected to reflect signal propagation dynamics along the axonal tree (*30, 31*).

**Table 2:** Morphological features of the axonal tree.

|  | Description |
| --- | --- |
| 1 | The number of branches |
| 2 | Symmetry – mean of the ratio between the number of children in each daughter branch |
| 3 | Maximum branch order |
| 4 | Mean branch order |
| 5 | Difference between maximum and minimum on the x-coordinates ($\mu m$) |
| 6 | Difference between maximum and minimum on the y-coordinates ($\mu m$) |
| 7 | Difference between maximum and minimum on the z-coordinates ($\mu m$) |
| 8 | Sholl analysis – the number of branch intersections at a radius of $100\mu m$ from the soma in 3D |
| 9 | Sholl analysis – the number of branch intersections at a radius of $200\mu m$ from the soma in 3D |
| 10 | Sholl analysis – the number of branch intersections at a radius of $300\mu m$ from the soma in 3D |
| 11 | The number of branches longer than $200\mu m$ |
| 12 | The number of branches longer than $300\mu m$ |
| 13 | The number of branches longer than $400\mu m$ |
| 14 | Maximum path length, the distance from the soma to the farther leaf ($\mu m$) |
| 15 | Minimum path length, the distance from the soma to the closer leaf ($\mu m$) |
| 16 | Mean path length, the distance from the soma to the closer leaf ($\mu m$) |
| 17 | Maximum branch length ($\mu m$) |
| 18 | Mean branch length ($\mu m$) |
| 19 | The total length of all branches in the axonal tree ($\mu m$) |
| 20 | Maximum branch length divided by the square root of the branch diameter |
| 21 | Mean branch length divided by the square root of the branch diameter |
| 22 | Maximum branch diameter ($\mu m$) |
| 23 | Mean branch diameter ($\mu m$) |
| 24 | The number of GRs above 2 |
| 25 | The number of GRs above 3 |
| 26 | The maximum GR value of all branching points |
| 27 | The mean GR value of all branching points |
| 28 | The percentage the of bifurcations with GR above 2 |

To avoid biases in classification due to unequal group sizes, and to decrease simulation time we downsampled our data to include 16 neurons in each group. These neurons were selected as the reconstructions with the highest number of diameter values measured from each group. We applied a 4-fold cross validation scheme with 1,000 repeats, and used a multinomial logistic regression approach with *l*^2^ regularization to classify the data. The resulting *F*_1_-scores are presented in Fig. 1H. *F*_1_-scores range between 0.702 for basket cells, and 0.902 for chandelier, resulting in an average *F*_1_-score of 0.777, based only on the axonal tree’s morphological parameters. These results are supported by the fact that chandelier cells are, indeed, the easiest cells for experts to classify manually (*5*). Furthermore, basket cells are heterogeneous, commonly divided into large, nest, and small basket cell subclasses (*26*). A noticeable similarity between the axonal trees of the double-bouquet and Martinotti cells can be seen in Fig. 1H. This resemblance supports previous studies that showed the similarity between the electrophysiological properties of these two types of neurons (*32*). A complement analysis of unsupervised clustering was performed on the morphological parameters for each neuron and is presented in fig. S1.

**Figure 1:**
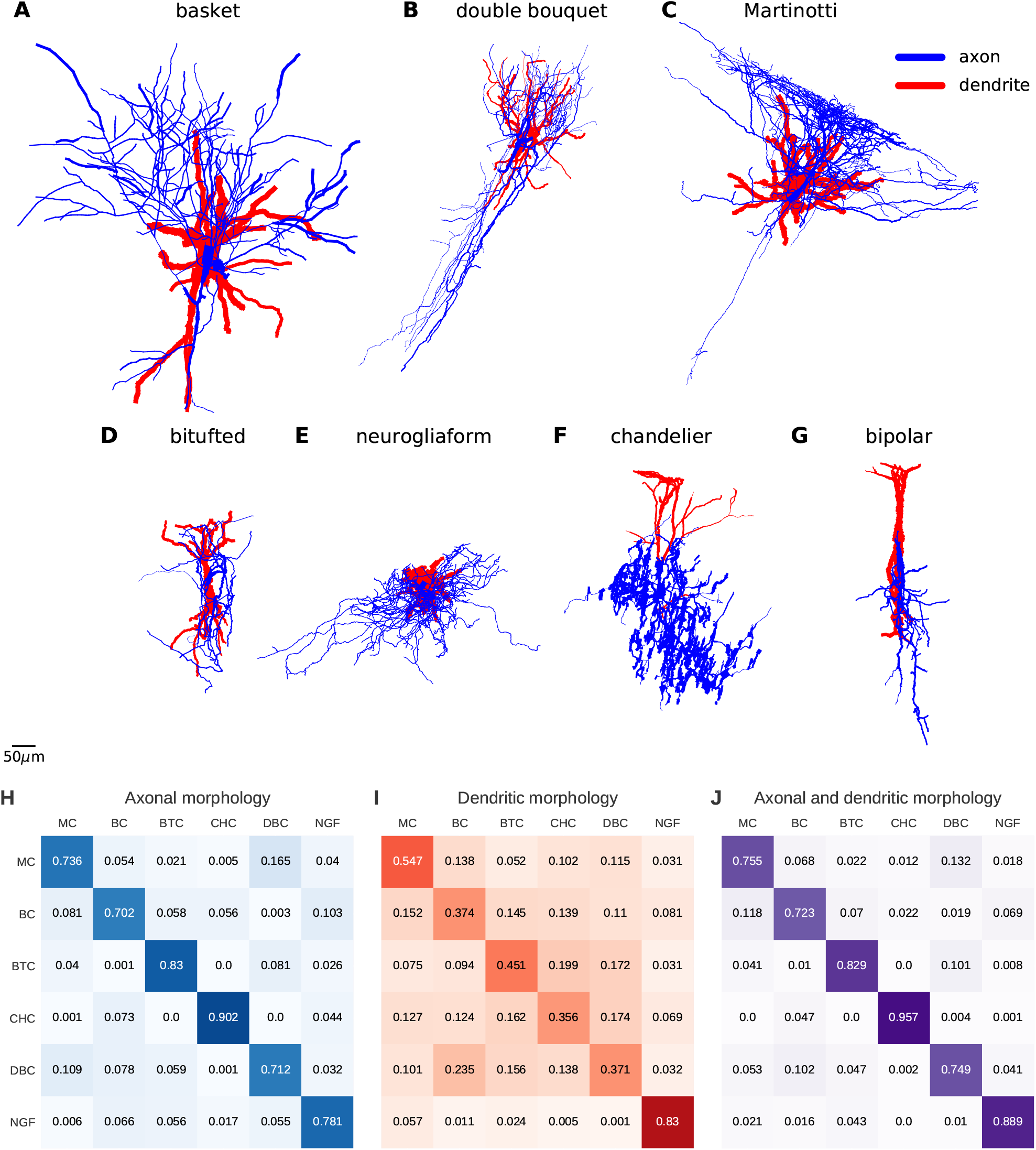
Classification by morphology. Representative examples of different interneuronal types. Line width is proportional to the axonal (blue) or dendritic (red) segment’s corresponding diameter. Data are projected into the XY plane. Cells used for visualizations are as follows: **A**. NMO_06143, **B**. NMO_61613, **C**. NMO_79459, **D**. NMO_ 61580, **E**. NMO_37062, **F**. NMO_04548, and **G**. NMO_61602. *F*_1_-score matrices for **H**. Axonal tree morphology only (average *F*_1_-score: 0.777), **I**. Dendritic tree morphology only (average *F*_1_-score: 0.488), and **J**. Axonal and dendritic tree morphologies combined (average *F*_1_-score: 0.817).

To compare the above classification based on axonal tree morphology to a more common classification based on dendritic tree morphology, we applied an analogous classification approach to the dendritic trees of the *same* neurons. The resulting *F*_1_-scores are presented in Fig. 1I. The dendritic tree-based classification better detects neurogliaform cells (*F*_1_-score of 0.83 compared to 0.356–0.547 in other cell types). This result agrees with the observation that neurogliaform cells are known for their thinness and abundance of radiating dendrites (*33*). In fact, of the six cell types, this is the only case in which the dendritic tree-based classification performs better than the axonal tree-based classification (*F*_1_-score of 0.83 compared to 0.781). Interestingly, the dendritic tree-based classification performs poorly on chandelier cells (*F*_1_-score of 0.356), in contrast to the axonal tree-based classification, that classifies these cells with a very high success rate (*F*_1_-score of 0.902). Note that bitufted cells are better differentiated by the axonal tree, even though their name was coined due to their dendritic tree structure.

We next combined axonal and dendritic tree morphology parameters and applied the same classification scheme as before. This resulted in an improved classification performance: the average *F*_1_-score increased from 0.777 for axonal trees only and 0.488 for dendritic trees only to 0.817 for the two combined (Fig. 1J).

The fitted classification models allow us to quantify the contribution of each feature to the classification (fig. S2). For example, neurogliaform cells are characterized by symmetrical topology, high Sholl values at 100*μm*, and high values of mean GR of the axonal tree. In the dendritic tree, however, they are characterized by low values of mean GR. In contrast, Martinotti cells have low values of the mean GR in the axonal tree. Bitufted cells have high values of mean branch length and mean branch length divided by the square root of diameter; Chandelier cells are characterized by high values of the maximum dendritic branch length and the maximum path length.

### Classification of interneuron types by signal propagation dynamics

To study signal propagation dynamics, we measured the response to current stimulus pulses injected into the soma at various frequencies along the axonal tree. Figure 2 shows an example of simulated neuronal activity along axonal branches of a basket cell. In this example, we used the membrane properties of the ‘continuous non-accommodating’ (cNAC) e-type, obtained from the BBP repertoire to simulate signal propagation. In the soma, all the stimulus pulses lead to action potential (denoted as ‘1’), and in the other probed locations intermitted trains occur (denoted as ‘2–4’).

**Figure 2:**
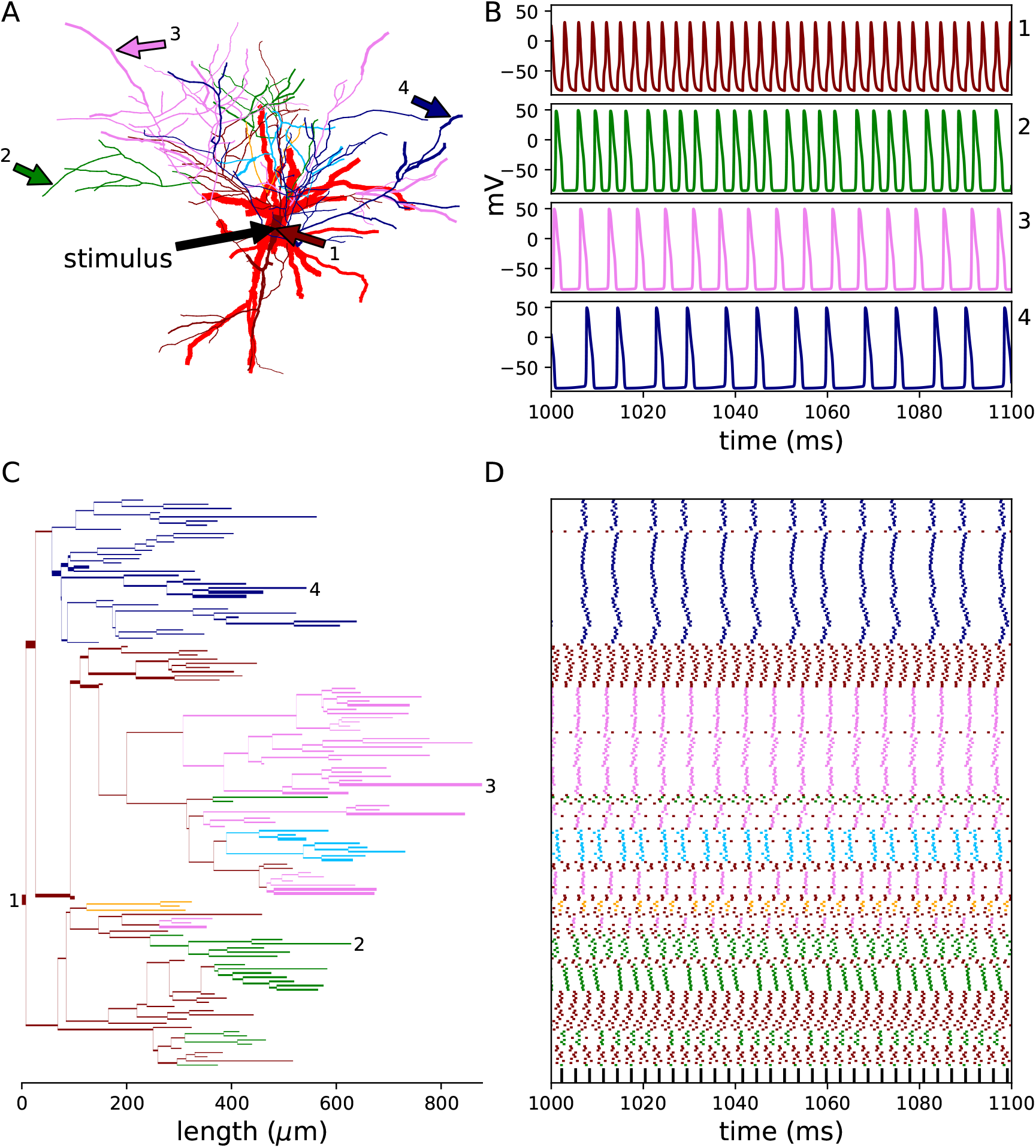
Activity recorded along the axonal tree. **A**. An XY projection for a basket cell (NMO_06143). The arrows indicate the location in which the stimulus is induced and the locations in which the propagated signal is recorded. Dendrites are in red, and the axonal branches are colored according to the firing pattern response. **B**. An example of firing patterns in four different locations. Panels 1–4 (indicated in the top right part of each panel) correspond to the arrows shown in (**A**). **C**. Axonogram: a dendrogram-like graph of the axonal tree only. Horizontal line widths indicate axonal diameters. **D**. Raster plot of the electrical activity; each row represents the activity at the corresponding (same height) axonal branch in (**C**). Line color indicates the fraction of spike train that propagates: maroon – 1, orange – 0.75, deep sky blue – 0.66, violet – 0.5, and navy – 0.375. Black squares on the bottom row indicate the current pulses applied to the soma (330*Hz*). The response to the first 1,000*ms* is not shown, to rule out the influence of the initial condition.

Figure 3 shows the electrical response along the axonal tree for different stimulus frequencies for the cNAC e-type. At 200*Hz* (Fig. 3A) the axonal tree is split into two subtrees, each exhibits a different firing pattern, in particular, an uninterrupted train and a ‘1:1’ pattern in which every other pulse propagates. For a stimulus frequency of 300*Hz* (Fig. 3B), there are 8 subtrees with three different response types, and at 400*Hz* (Fig. 3C), there are 23 subtrees with five different response types. We counted the number of subtrees as a function of stimulus frequency (Figure 3D), and obtained a characteristic curve that will be used for the classification. In addition to the cNAC, we generated three other curves from the cAC, bNAC, and bAC e-types. The decision to focus on these e-types resulted from an analysis that showed a high degree of propagating signal similarity between e-types (see fig. S3).

**Figure 3:**
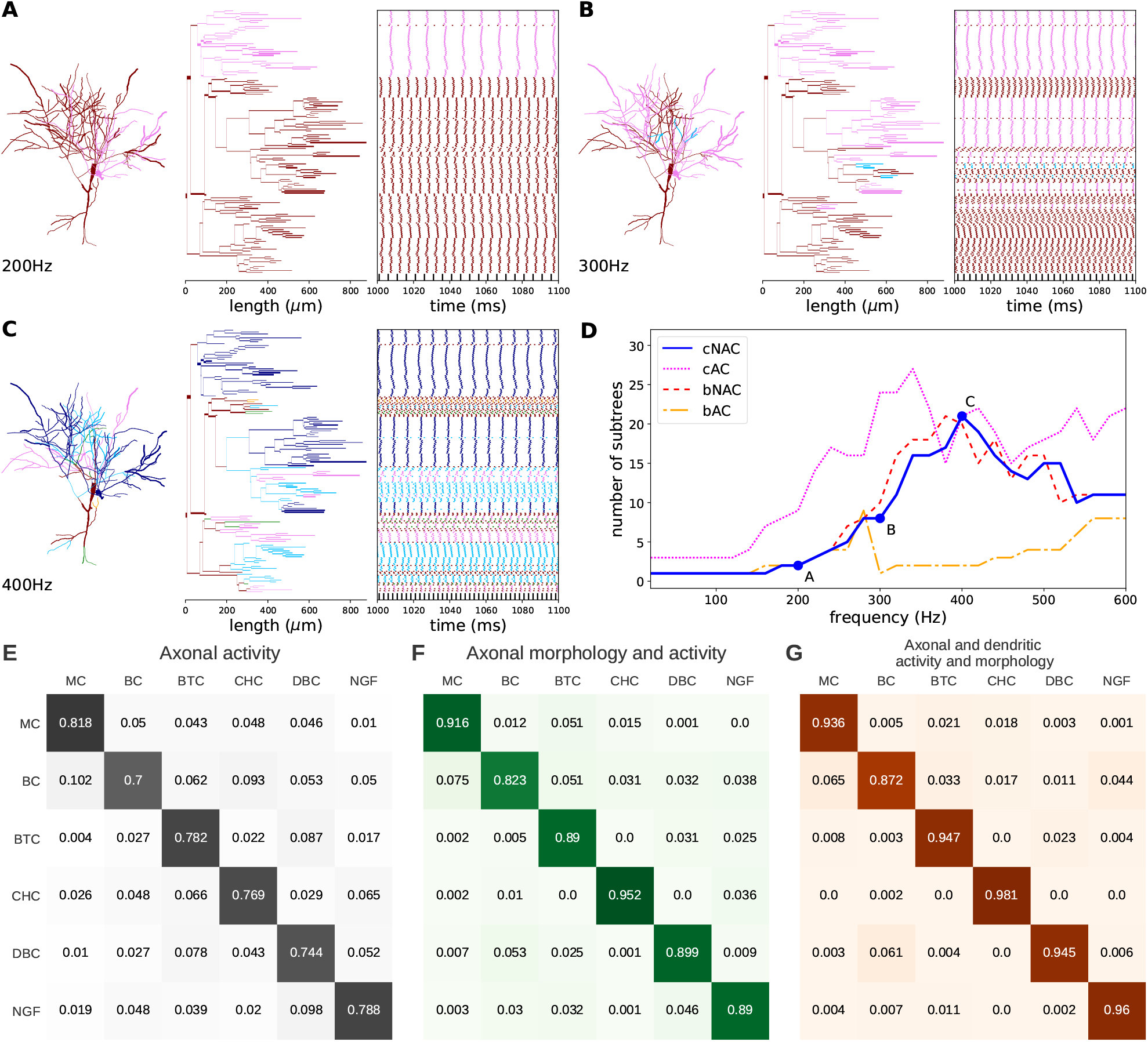
Effects of stimulus frequency on the signal propagation dynamics. Axonograms showing the responses of the same interneuron (as in Fig. 2) to three stimulus frequencies are presented in **A**. (200*Hz*), **B**. (300*Hz*), and **C**. (400*Hz*). **D**. The number of subtrees as a function of stimulus frequency. The dots (marked in ‘A’, ‘B’, and ‘C’) correspond to the scenarios presented in (**A**), (**B**), and (**C**). *F*_1_-score matrices for **E**. Activity only (average *F*_1_-score: 0.767), **F**. Axonal morphology and activity combined (average *F*_1_-score: 0.895), and **G**. Axonal and dendritic morphology and activity combined (average *F*_1_-score: 0.94).

We then tested the possibility of using neuronal activity signatures to classify interneuron types. To this end, we have engineered activity-based features from Hill diversity indexes (see Methods). Figure S4 shows the mean and standard deviation of the *q* = 0 diversity index as a function of the stimulus frequency. Note that for *q* = 0, the diversity index is equal to the number of subtrees. Similar to the morphology-based classification, we used multinomial logistic regression, and assessed it with a 4-fold cross validation scheme with 1,000 repeats. Figures 3E presents the classification results based on the axonal tree activity (an average *F*_1_-score of 0.767). Combining axonal morphology (Fig. 1H) with axonal tree activity (Fig. 3E) improves the classification’s average *F*_1_-score from 0.777 and 0.767 to 0.895 (Fig. 3F). Including the morphology of the dendritic tree as well resulted in an average *F*_1_-score of 0.94 (Fig 3G). The classification model results for each neuron are presented in Table S1.

## Discussion

Numerous studies published in recent years used a variety of approaches to classify neurons. Standard classification approaches are based on morphology. In particular, they utilize soma position, dendritic geometry, general axonal projections, and connectivity features (*4*). Here we used the axonal tree morphology for interneuron classification, resulting in better results compared with classification based on the dendritic tree morphology. Combining axonal tree morphology with dendritic tree morphology and activity patterns further improved the classification results. For the data analyzed here, the average *F*_1_-score changed from 0.777 for axonal trees and 0.488 for dendritic trees to 0.817 for the two combined. Combining also the activity patterns significantly improved the *F*_1_-score to 0.94.

Interneuron subtypes are known to shape the electrophysiological activity dynamics (*34*). Therefore, the use of activity-based features as classifiers is important (*35*). Teeter *et al*. used simple generalized leaky integrate-and-fire point neuron models to classify transgenic lines (*36*). Previous studies have also shown how dendritic tree geometry affects the electrical activity in neurons. It was demonstrated that in activity simulations taking into account the dendritic tree morphology, rather than a point neuron, can capture local non-linear effects (*37*). In this study, we used realistic neuronal morphologies and dynamics for classification of interneurons. To this end, the soma was stimulated with a wide range of current pulses frequencies, since it was shown that high-frequency trains may exist in fast-spiking neurons and in bursting (*38, 39*). Then, we recorded the simulated activity along each branch of the axonal tree, and not only at the soma. This revealed diverse response patterns already at the single neuron level. Importantly, we showed that these firing patterns can be used to classify interneurons into their known subtypes. Our results indicate that it is beneficial for modeling signal propagation dynamics to include the full biophysical neuronal structure including the axonal tree. In particular, precise measurements of neuronal processes were shown here to strongly affect simulation results in single neuron models. In the future, advanced imaging tools may be used to explore the activity along the axonal tree in real neurons. This will allow to study the effects of geometry on the accuracy of activity simulations.

Due to the lack of accepted nomenclature and no complete agreement between experts, the labels given by a specific lab, provided by *NeuroMorpho.Org*, are prone to errors. It can be seen in table S1 and in fig. S1, that there are neurons that evidently belong to a different type than the one tagged by *NeuroMorpho.Org* (e.g., NMO_37137 and NMO_61618). Discarding these neurons may further improve classification results. Despite tremendous progress in imaging and reconstruction techniques, the amount of quality data is not sufficient. More data of high-resolution reconstructions from diverse sources are of utmost importance for more comprehensive species dependent neuron classification. When such additional data become available, neurons of rats, mice, and humans, from different brain regions and layers, could be independently classified. The electrical membrane properties of the reconstructions used here, were fitted by the BBP to the soma, dendrites, and the axon initial segment, and do not include the entire axonal tree, axonal boutons, and myelin sheath (*26*). Nevertheless, these properties resemble a close approximation of an actual mechanism, and yielded an excellent classifications. Fitting the electrical membrane properties along all the axonal tree, can further improve our understanding of signal propagation in neurons.

Classifying neurons into subtypes is debatable; it is not yet clear that neurons can be described by a set of distinct classes or should they be treated as a continuum of phenotypes (*18*). The approach used here of combining dendritic and axonal tree morphologies with activity patterns can be utilized in an unsupervised fashion to examine the clusters of subtypes without prior assumptions. This investigation will also advance standardization toward consensus regarding neuronal type nomenclature.

## Materials and Methods

### Simulations

Digitally reconstructed neurons were downloaded from *NeuroMorpho.Org* version 7.4 (*28*). All reconstructions were produced in the same staining method, by bright-field images of biocytin filled neurons. Each neuron’s reconstructed data is stored in an SWC file. We used the notation of branch to describe an axonal section between two branching points, or between a branching point and a termination point (leaf), and a segment to describe a small compartment in 3D space. Several successive segments were used to construct a branch. The SWC files were imported into *NEURON* simulation using the *Import3D* tool, which converts all the segments of each branch into equivalent diameter cables. The neuronal activity simulations were conducted using *NEURON* simulation environment version 7.5 embedded in Python 2.7.13 (*40*). The same version of Python was used for all other analyses presented here.

### Membrane electrical properties

To simulate signal propagation dynamics, ion channel mechanisms with different densities were introduced into the reconstructed neurons. For realistic modeling, we used membrane properties borrowed from the BBP (*26*). These e-types were fitted to experiments produced in the cortex neurons of Wistar (Han) rats at a temperature of 34°*C*. Each e-type is constructed from specific ion channel types with varying densities at the soma, axons, and basal and apical dendrites. Details of the ion channels, their kinetics, and other parameters can be found in the NMC portal (*25*) and in the attached files there. The specific equations and parameters used here were taken from the following reconstructions: L23_LBC_cNAC187_5, L23_DBC_cACint209_1, L23_LBC_bNAC219_1, L5_LBC_bAC217_4, and L5_STPC_cADpyr.

Current pulses were stimulated in the soma, with an amplitude of 20*μA* and a duration of 1*ms*, for a range of frequencies. Electrical responses were recorded at the center of each axonal branch. For the raster plots (e.g., Fig. 2D), a spike was defined when the voltage peak amplitude exceeded a zero voltage threshold. Voltage peaks that were separated by less than 1*ms* were discarded to avoid discretization errors.

### Classification

Supervised classification was performed using multinomial logistic regression. We used the *LogisticRegression* function from the *Scikit-learn* python library, with an *l*^2^ regularization penalty. For the classification, 1,000 4-fold cross validation repeats were produced, i.e., 1,000 choices of 75% of the data for training and 25% for testing the model. All morphological parameters were first log-transformed, and then standardized for this classification. The parameters for both the training and test sets were standardized according to the mean and standard deviation of the training set. Feature selection was performed in a recursive way where in each iteration, the worst feature of each neuronal type was discarded, to achieve the optimize features for the final model. Sensitivity was calculated by normalizing each value in the confusion matrix by the sum of the row to which it belongs. Precision was calculated similarly but normalization was done according to the column of the confusion matrix. *F*_1_-score is defined as the harmonic average of sensitivity and precision (Equation 1).

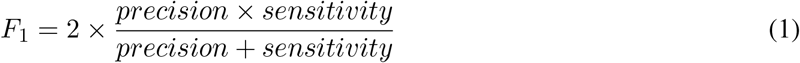

For activity-based classification, the Hill diversity index (Equation 2) was calculated as a function of frequency for each interneuron type. The Hill diversity index was calculated for *q* ∈ {0, 1}, for four e-types, and two definitions of a subgroup: 1. A subtree with an identical response, and 2. A set of branches with an identical response (possibly in more than one subtree). The Hill diversity index is defined as

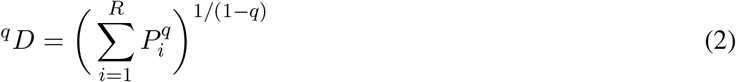

where *R* is the number of groups (“species” according to its original definition), and *P_i_* is the normalized number of members in each group. ^0^*D* equals the number of groups, and ^1^*D* converges to the exponent of Shanon Entropy (Equation 3).

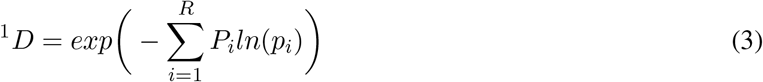

### Code availability

The code for the models and simulations is publicly available on Github: https://github.com/NetanelOfer/Axonal_tree_classification.

## Supplementary Materials

Table S1. Detailed confusion matrix.

Fig. S1. A heatmap comparing the distributions of morphological parameters.

Fig. S2. Logistic regression coefficients.

Fig. S3. Comparison between 10 e-types from the BBP.

Fig. S4. Axonal response characterized by diversity index as a function of stimulus frequency.

## Supplementary material

**Table S1:** Detailed confusion matrix. The results of the classification based on the axonal and dendritic morphology and activity (an elaboration of Fig. 3G). The maximum number in each row is indicated by blue background.

| Type | NMO | MC | BC | BTC | CHC | DBC | NGF |
| --- | --- | --- | --- | --- | --- | --- | --- |
| Martinotti | NMO_06140 | 250 | 4 | 0 | 0 | 0 | 0 |
|  | NMO_36965 | 194 | 0 | 0 | 56 | 1 | 0 |
|  | NMO_37190 | 245 | 2 | 2 | 0 | 3 | 1 |
|  | NMO_37290 | 202 | 0 | 38 | 17 | 0 | 0 |
|  | NMO_37291 | 256 | 0 | 0 | 0 | 0 | 0 |
|  | NMO_37297 | 235 | 0 | 0 | 0 | 0 | 0 |
|  | NMO_37301 | 233 | 5 | 0 | 0 | 0 | 0 |
|  | NMO_37672 | 269 | 0 | 0 | 0 | 0 | 0 |
|  | NMO_37787 | 201 | 3 | 47 | 0 | 0 | 2 |
|  | NMO_79457 | 213 | 0 | 0 | 0 | 0 | 0 |
|  | NMO_79458 | 269 | 0 | 0 | 0 | 0 | 0 |
|  | NMO_79459 | 245 | 0 | 0 | 0 | 0 | 0 |
|  | NMO_79462 | 270 | 4 | 0 | 0 | 0 | 0 |
|  | NMO_79463 | 230 | 0 | 0 | 0 | 8 | 0 |
|  | NMO_79464 | 236 | 0 | 0 | 0 | 0 | 0 |
|  | NMO_79465 | 260 | 0 | 0 | 0 | 0 | 0 |
| basket | NMO_06143 | 1 | 236 | 0 | 0 | 0 | 0 |
|  | NMO_37015 | 0 | 268 | 0 | 5 | 0 | 0 |
|  | NMO_37112 | 2 | 227 | 0 | 1 | 5 | 0 |
|  | NMO_37117 | 129 | 108 | 2 | 7 | 2 | 9 |
|  | NMO_37137 | 0 | 73 | 0 | 30 | 4 | 151 |
|  | NMO_37188 | 0 | 203 | 30 | 0 | 15 | 0 |
|  | NMO_37310 | 0 | 254 | 0 | 0 | 0 | 0 |
|  | NMO_37311 | 2 | 287 | 0 | 1 | 0 | 0 |
|  | NMO_37504 | 0 | 248 | 0 | 0 | 0 | 1 |
|  | NMO_37505 | 0 | 232 | 0 | 0 | 1 | 15 |
|  | NMO_37529 | 15 | 207 | 0 | 24 | 0 | 0 |
|  | NMO_37841 | 0 | 232 | 0 | 0 | 1 | 0 |
|  | NMO_37844 | 0 | 148 | 101 | 0 | 7 | 0 |
|  | NMO_37862 | 2 | 260 | 0 | 0 | 6 | 1 |
|  | NMO_79468 | 96 | 159 | 0 | 0 | 1 | 0 |
|  | NMO_79469 | 17 | 212 | 0 | 0 | 0 | 0 |
| bitufted | NMO_37096 | 0 | 0 | 250 | 0 | 0 | 0 |
|  | NMO_37099 | 0 | 0 | 254 | 0 | 0 | 0 |
|  | NMO_37110 | 0 | 0 | 256 | 0 | 0 | 0 |
|  | NMO_37127 | 0 | 0 | 255 | 0 | 2 | 0 |
|  | NMO_37244 | 0 | 3 | 260 | 0 | 0 | 0 |
|  | NMO_37296 | 0 | 0 | 254 | 0 | 0 | 0 |
|  | NMO_37302 | 0 | 0 | 243 | 0 | 0 | 0 |
|  | NMO_37316 | 0 | 0 | 252 | 0 | 0 | 0 |
|  | NMO_37324 | 0 | 0 | 244 | 0 | 0 | 0 |
|  | NMO_37385 | 0 | 0 | 267 | 0 | 0 | 0 |
|  | NMO_37691 | 0 | 0 | 241 | 0 | 1 | 0 |
|  | NMO_37704 | 0 | 0 | 260 | 0 | 0 | 0 |
|  | NMO_37776 | 0 | 0 | 236 | 0 | 0 | 0 |
|  | NMO_37781 | 0 | 0 | 226 | 0 | 0 | 0 |
|  | NMO_61570 | 34 | 4 | 198 | 0 | 3 | 0 |
|  | NMO_61580 | 0 | 5 | 152 | 0 | 84 | 17 |
| chandelier | NMO_04548 | 0 | 0 | 0 | 241 | 0 | 0 |
|  | NMO_07472 | 0 | 0 | 0 | 249 | 0 | 0 |
|  | NMO_35831 | 0 | 0 | 0 | 248 | 0 | 0 |
|  | NMO_36983 | 0 | 0 | 0 | 261 | 0 | 0 |
|  | NMO_37114 | 1 | 0 | 0 | 245 | 1 | 0 |
|  | NMO_37133 | 0 | 4 | 0 | 249 | 0 | 0 |
|  | NMO_37138 | 0 | 0 | 0 | 264 | 0 | 0 |
|  | NMO_37247 | 0 | 1 | 0 | 279 | 0 | 0 |
|  | NMO_37391 | 0 | 0 | 0 | 244 | 0 | 0 |
|  | NMO_37424 | 0 | 0 | 0 | 235 | 0 | 0 |
|  | NMO_37537 | 0 | 0 | 0 | 255 | 0 | 0 |
|  | NMO_37548 | 0 | 4 | 0 | 240 | 0 | 0 |
|  | NMO_37763 | 0 | 0 | 0 | 236 | 0 | 0 |
|  | NMO_37764 | 0 | 0 | 0 | 240 | 0 | 0 |
|  | NMO_37818 | 0 | 0 | 0 | 245 | 0 | 0 |
|  | NMO_37821 | 0 | 0 | 0 | 261 | 0 | 0 |
| double-bouquet | NMO_37148 | 0 | 0 | 3 | 0 | 264 | 6 |
|  | NMO_37171 | 0 | 0 | 5 | 0 | 237 | 0 |
|  | NMO_37178 | 0 | 0 | 0 | 1 | 252 | 0 |
|  | NMO_37184 | 0 | 0 | 1 | 0 | 240 | 0 |
|  | NMO_37363 | 0 | 0 | 0 | 0 | 248 | 0 |
|  | NMO_37422 | 1 | 0 | 0 | 0 | 256 | 0 |
|  | NMO_37423 | 0 | 0 | 0 | 0 | 243 | 0 |
|  | NMO_37469 | 0 | 0 | 0 | 0 | 260 | 0 |
|  | NMO_37475 | 0 | 30 | 0 | 1 | 228 | 0 |
|  | NMO_37694 | 0 | 0 | 0 | 0 | 264 | 0 |
|  | NMO_37695 | 0 | 0 | 1 | 0 | 263 | 0 |
|  | NMO_37699 | 0 | 0 | 0 | 0 | 263 | 0 |
|  | NMO_37724 | 0 | 1 | 2 | 0 | 245 | 17 |
|  | NMO_37762 | 0 | 0 | 0 | 0 | 248 | 0 |
|  | NMO_61613 | 3 | 42 | 0 | 0 | 207 | 0 |
|  | NMO_61618 | 9 | 161 | 3 | 0 | 63 | 3 |
| neurogliaform | NMO_06138 | 0 | 0 | 0 | 0 | 0 | 244 |
|  | NMO_06139 | 14 | 23 | 22 | 0 | 0 | 191 |
|  | NMO_37059 | 0 | 0 | 20 | 0 | 0 | 222 |
|  | NMO_37061 | 0 | 0 | 0 | 0 | 0 | 271 |
|  | NMO_37062 | 0 | 0 | 0 | 0 | 0 | 244 |
|  | NMO_37063 | 0 | 0 | 0 | 0 | 0 | 252 |
|  | NMO_37075 | 0 | 0 | 0 | 0 | 0 | 266 |
|  | NMO_37076 | 0 | 0 | 0 | 0 | 0 | 230 |
|  | NMO_37243 | 0 | 0 | 0 | 0 | 0 | 234 |
|  | NMO_37517 | 0 | 0 | 0 | 0 | 0 | 237 |
|  | NMO_37623 | 0 | 0 | 0 | 0 | 2 | 273 |
|  | NMO_37625 | 0 | 0 | 0 | 0 | 0 | 244 |
|  | NMO_37640 | 0 | 0 | 3 | 0 | 0 | 250 |
|  | NMO_37661 | 0 | 0 | 0 | 0 | 7 | 251 |
|  | NMO_37720 | 1 | 2 | 1 | 1 | 0 | 226 |
|  | NMO_37721 | 0 | 3 | 0 | 0 | 0 | 240 |

**Figure S1:**
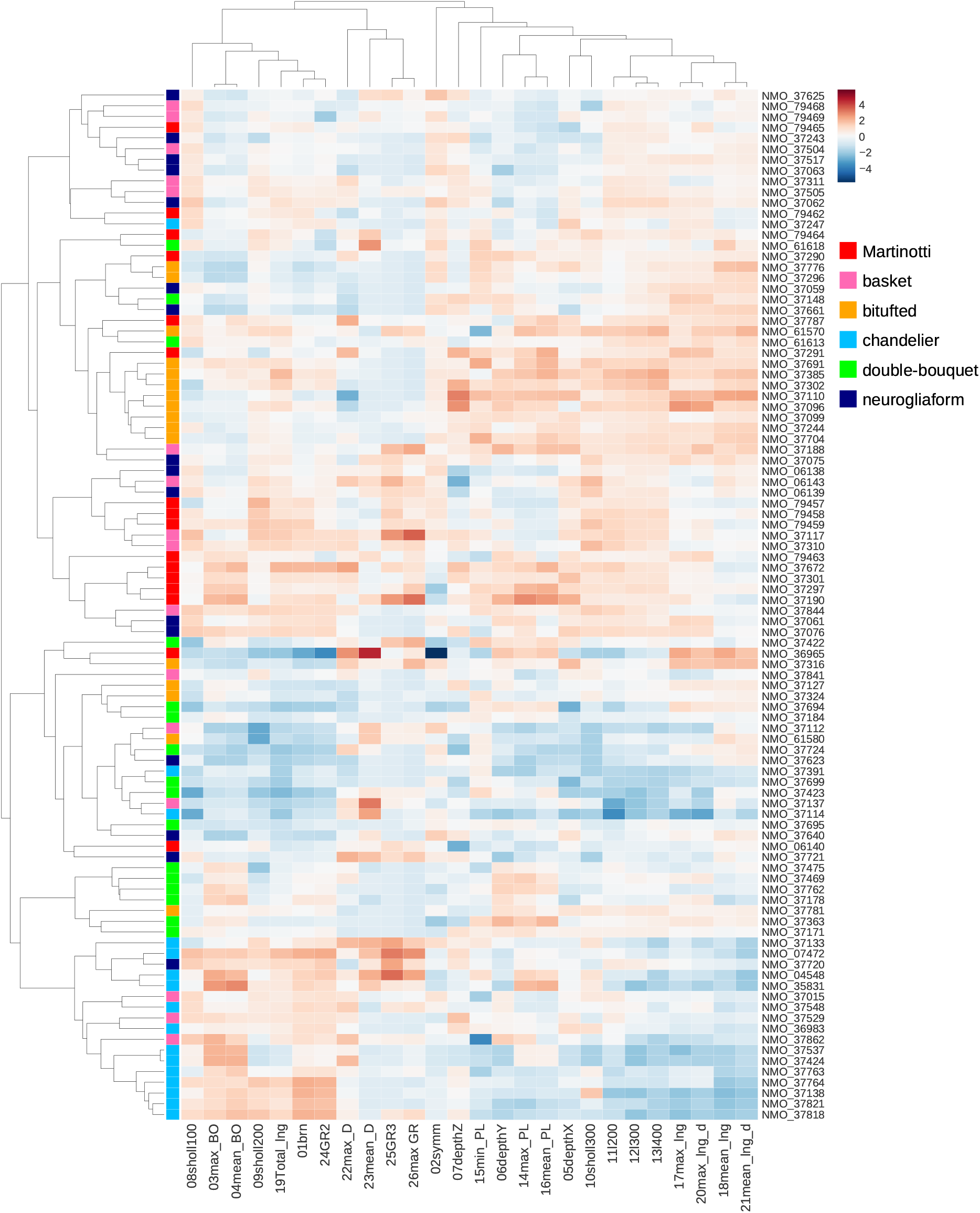
A heatmap comparing the distributions of morphological parameters. The parameters were transferred to *z*-score values. The *NeuroMorpho.Org* IDs are indicated for each neuron reconstruction on the right side, and the interneuron subtype is indicated by the ‘row color’ on the left side. The columns are the 28 morphological parameters organized according to the above dendrogram. The left dendrogram clusters the neurons in an unsupervised manner.

**Figure S2:**
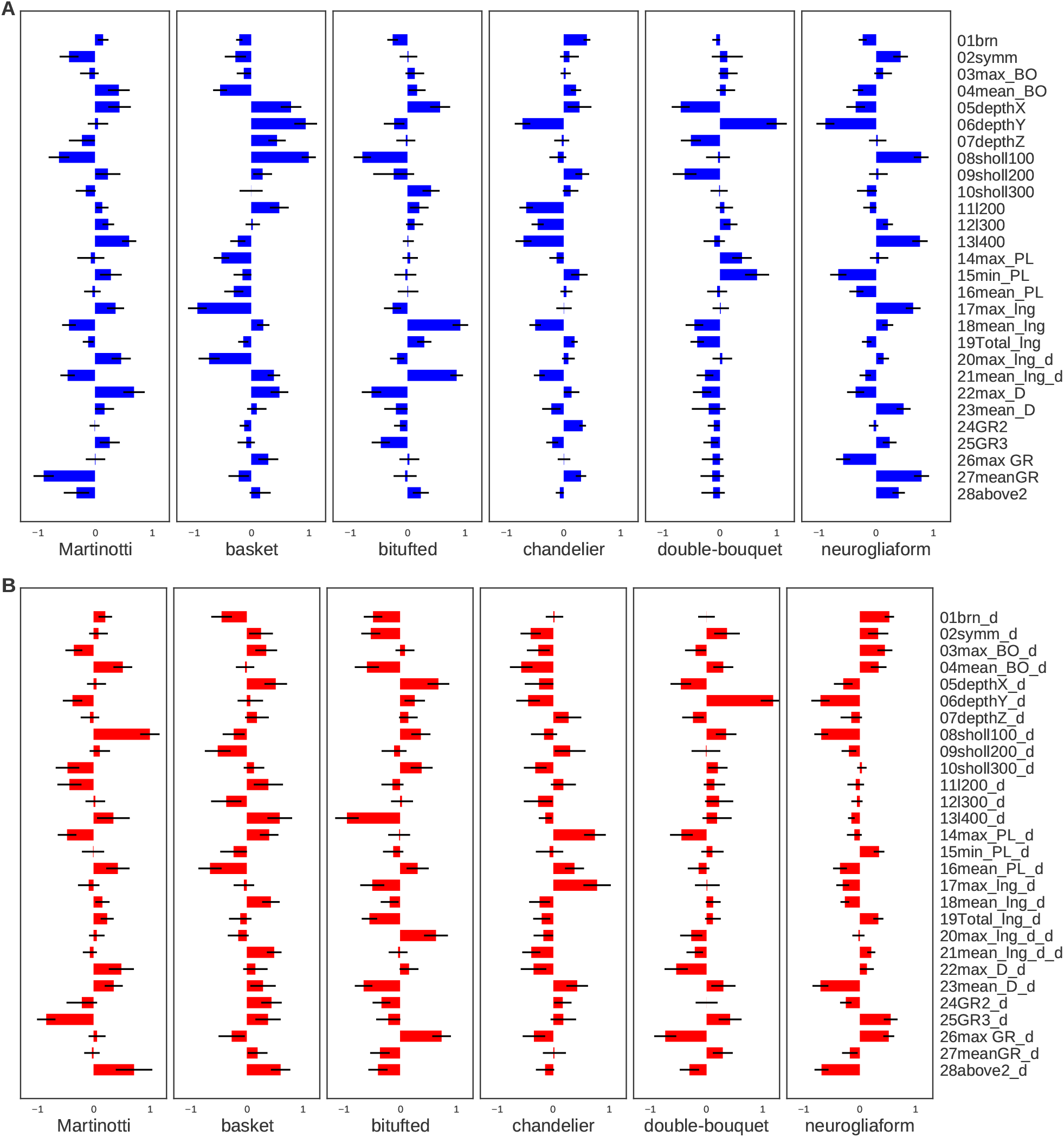
Logistic regression coefficients. The average of the logistic regression coefficients for the 1,000 repeats of the axonal (**A**) and dendritic (**B**) trees’ morphology. Positive values indicate the significance of this feature for the specific interneuron type.

**Figure S3:**
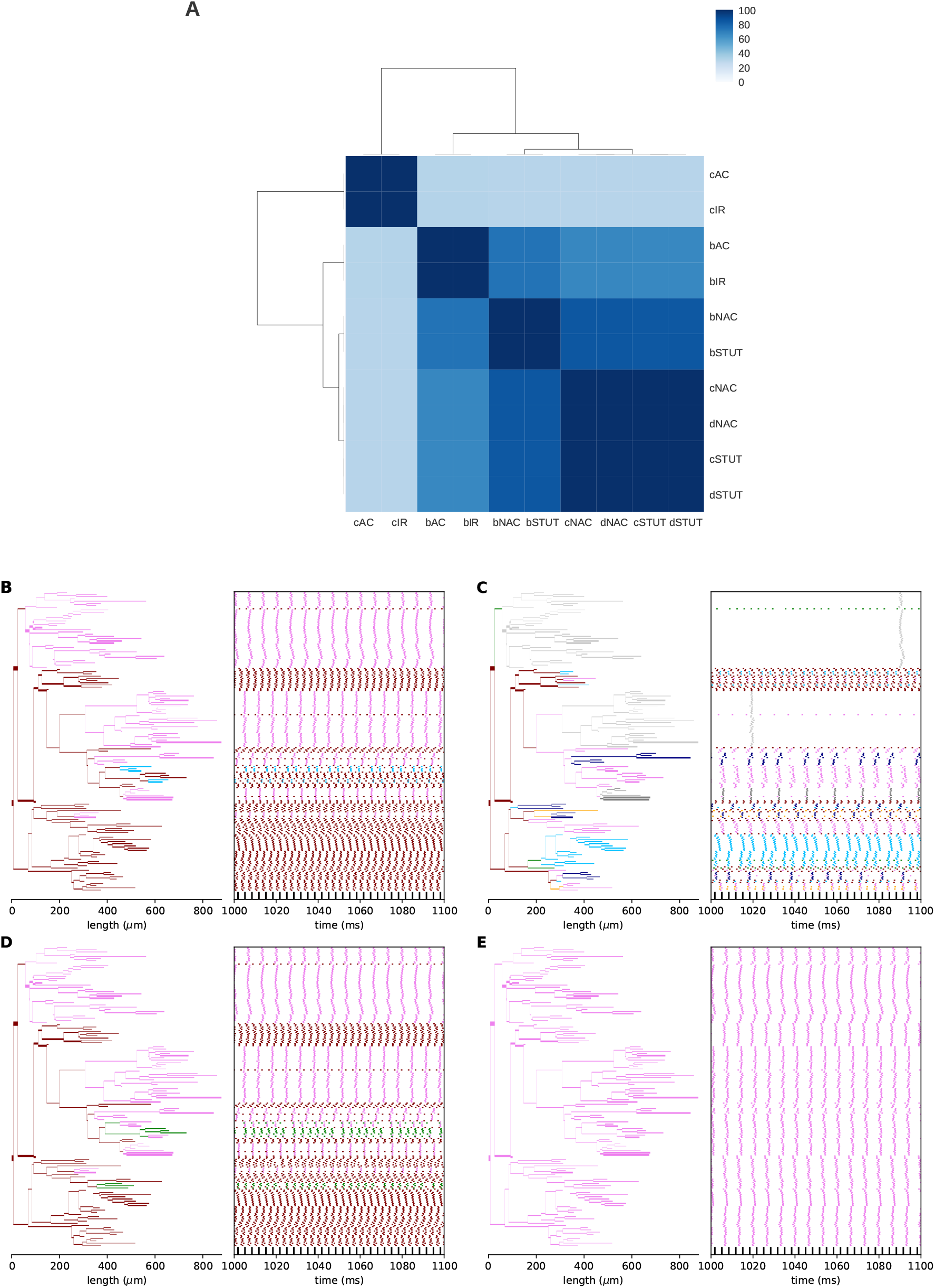
Comparison between 10 e-types from the BBP. A heatmap showing the similarity between all 10 interneuron e-types under current pulse frequencies of 100*Hz*, 200*Hz*, 300*Hz*, and 400*Hz* in terms of firing pattern, in a basket cell (NMO_06143). In the original experiments conducted by BBP, an elongated current step was induced, resulting in significant differences between these 10 e-types. In our case, the soma was stimulated with strong current pulses, leading to very similar responses in several e-types. Hence, we chose to focus on four e-types: cAC, bAC, bNAC, and cNAC. The response for stimulus frequency of 300*Hz*. **B**. ‘cNAC’ (the same as in Fig. 3B), **C**. ‘cAC’, **D**. ‘bNAC’, and **E**. ‘bAC’ e-types. The line color indicates the fraction of spike train that propagates: maroon – 1, orange – 0.75, deep sky blue – 0.66, violet – 0.5, navy – 0.375, dim gray – 0.2, and silver – 0. The same neuron as in Fig. 2.

**Figure S4:**
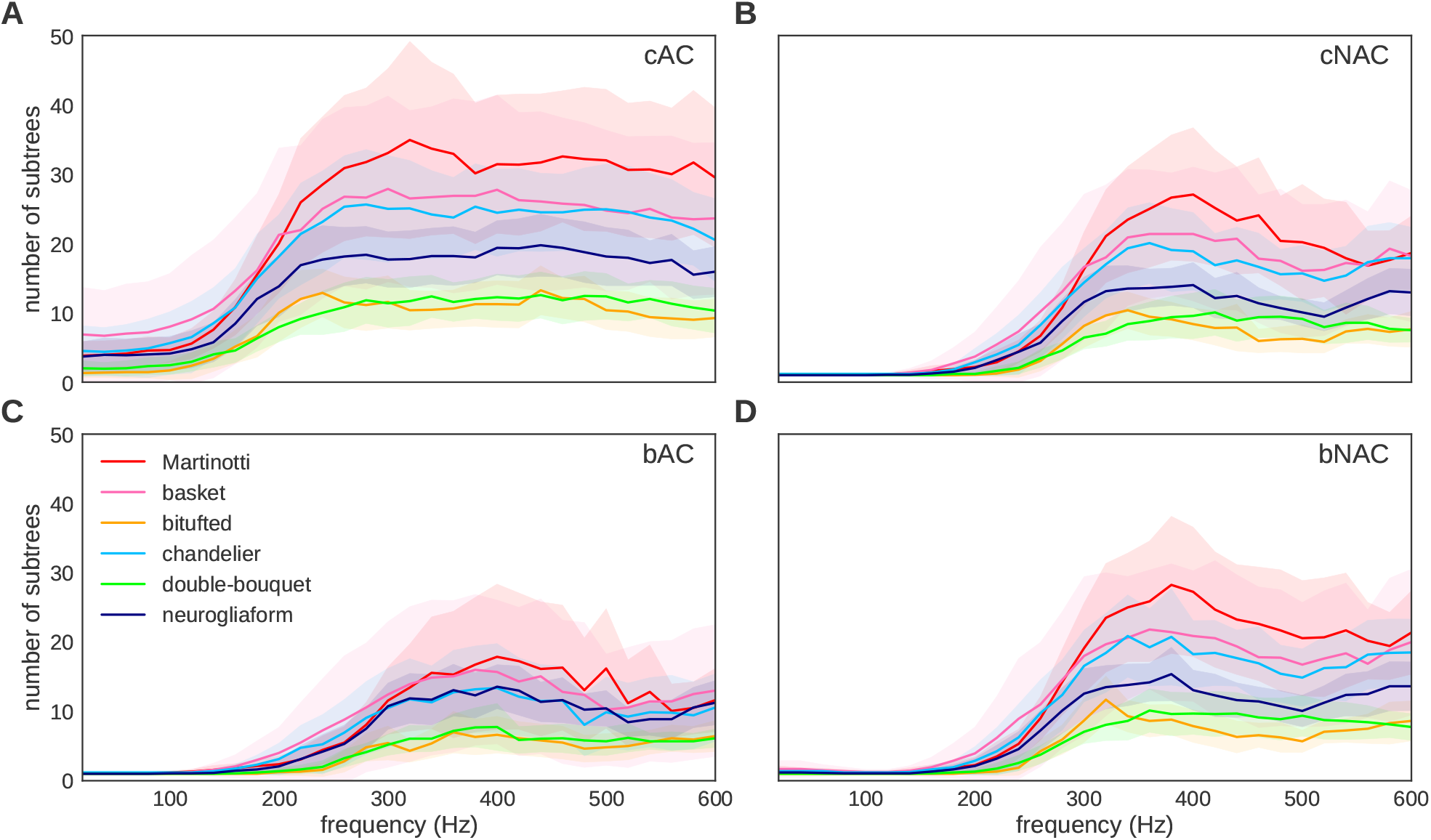
Axonal response characterized by diversity index as a function of stimulus frequency. Solid lines represent the mean diversity index (*q* = 0) and the shaded regions represent plus minus one standard deviation. Each panel shows the response for another e-type: **A**. Continuous accommodating (cAC), **B**. Continuous non-accommodating (cNAC), **C**. Burst accommodating (bAC), and **D**. Burst non-accommodating (bNAC).

